# Fast & Scrupulous: Gesture-Based Alarms Improve Accuracy and Reaction Times Under Various Mental Workload Levels. An ERSP Study

**DOI:** 10.1101/2022.03.21.485187

**Authors:** Eve Floriane Fabre, Bertille Somon, Valeria Baragona, Quentin Uhl, Mickaël Causse

**Author notes:** Correspondence: Eve Floriane Fabre.

## Abstract

In high-risk environments, fast and accurate responses to warning systems are essential to efficiently handle emergency situations. The aim of the present study was twofold: 1) investigating whether hand action videos (i.e., gesture alarms) trigger faster and more accurate responses than text alarm messages (i.e., written alarms), especially when mental workload (MWL) is high; and 2) investigating the brain activity in response to both types of alarms as a function of MWL. Regardless of MWL, participants (N = 28) were found to be both faster and more accurate when responding to gesture alarms than to written alarms. Brain electrophysiological results suggest that this greater efficiency might be due to a facilitation of the action execution, reflected by the decrease in mu and beta power observed around the response time window. These results suggest that gesture alarms may improve operators’ performances in emergency situations.

## **1.** INTRODUCTION

### 1.1 Reacting to emergency situations

In high-risk environments, such as transportation or nuclear power plants, fast and accurate reactions are essential to efficiently handle emergency situations and prevent potential catastrophic events. Since alarm systems play a critical role in triggering timely and appropriate responses in operators, particular care must be taken to ensure that their design maximizes the efficacy of the operator’s response in the event of an emergency (e.g., <u>Brown et al., 2000</u>; <u>Torralba et al., 2007</u>; <u>Wu & Li, 2018</u>).

Alarm systems are expected to distract operators from the task currently at hand and provide them with crucial information such as: 1) the gravity of the failure, 2) its cause, 3) its location and even sometimes 4) how to respond to it (<u>Guillaume, 2011</u>). Because stimulating two or more sensory channels of information (e.g., visual, auditory and/or haptic) facilitates both the detection and the reaction to alarms, emergency alarm systems are almost always multimodal usually composed of an auditory component and a visual component (e.g., <u>Alirezaee et al., 2017</u>; <u>Chan & Chan, 2006</u>; <u>Child & Wendt, 1938</u>; <u>Doyle & Snowden, 2001</u>; <u>Hughes et al., 1994</u>; <u>Liu, 2001</u>; <u>Schröger & Widmann, 1998</u>). On the one hand, auditory alarms are very effective to capture operators’ attention because they are omnidirectional and can be detected without having to gaze a specific area. On the other hand, visual alarms allow to provide more information to operators compared to auditory alarms (<u>Guillaume, 2011</u>; <u>Stevens et al., 2009</u>).

Emergency alarm systems generally combine symbolic (e.g., flashing lights, blares, or bips) and speech alarms (<u>Guillaume, 2011</u>). Symbolic alarms are very effective to attract the operator’s attention, even though their use implies a thorough and repeated training of operators to create and sufficiently consolidate the association between a symbolic alarm and the particular event it signals to guarantee satisfactory levels of retention and detection (e.g., <u>Momtahan, et al., 1993</u>; <u>Stevens et al., 2009</u>). While compared to symbolic alarms, speech alarms tend to trigger slower reactions in operators (e.g., <u>Wheale, 1982</u>), they allow to convey complex information (<u>Whittemore & Woods, 2021)</u>. They can either be *simple awareness warnings*, such as the Stall alarm, that indicate the nature of the emergency (e.g., in this case that the aircraft is about to enter a stall), but not how to respond to it (i.e., decreasing thrust and performing a nose-down pitch action; <u>IATA, 2018a</u>; <u>Whittemore & Woods, 2021</u>) or *pro- active instructions*, such as the Ground Proximity Warning System (GPWS; <u>EASA, 2007</u>), that indicate how to respond to the emergency (i.e., increasing thrust and performing a nose-up pitch action called “Pull Up”) without explicitly stating its nature (i.e., the aircraft is about to hit the ground; <u>IATA, 2018b; Moroze & Snow, 1999</u>).

### 1.2 Why operators sometimes fail to appropriately react to emergency situations

Despite the important efforts that have been made to improve emergency alarm systems, failures to effectively respond to emergency alarms are still responsible for a significant number of accidents (<u>Bliss, 2003</u>; <u>Kelly & Efthymiou, 2019</u>; <u>Scoot, 1996; Whittemore & Woods, 2021</u>). For instance, pilots’ delayed or inadequate reactions to the Stall or the GPSW alarms were found to be an important contributive factor in a significant part of respectively Controlled Flight Into Stall (CFIS; <u>IATA, 2018a; Oliver et al., 2019; Sherry & Mauro, 2014)</u> and Controlled Flight Into Terrain (CFIT; <u>Cooper, 1995</u>; <u>Moroze & Snow, 1999</u>; <u>Shappell & Wiegmann, 2001</u>; <u>2003</u>) accidents (i.e., when an airworthy aircraft under the complete control of the crew is inadvertently flown into respectively the ground or a stall), accounting for many fatalities in commercial aviation (<u>IATA, 2021</u>; <u>ICAO, 2019</u>).

Ergonomics flaws are one of the reasons why operators do not always properly respond to emergency alarms (<u>Scoot, 1996; Whittemore & Woods, 2021</u>). For instance, symbolic auditory alarms become hard to recognize for operators when many of them are played concurrently (e.g., <u>Causse et al., 2022</u>; <u>Momtahan, et al., 1993</u>), and speech auditory alarms can blend with operators’ communications and not be detected or properly understood (<u>Guillaume, 2011</u>). Moreover, visual alarms can be difficult to detect due to their small size and the important amount of concurrent visual information (<u>Corwin, 1995</u>).

Mental workload (MWL) can also affect operators’ response to emergency alarms (e.g., <u>Bliss & Dunn, 2000</u>; <u>Bliss, 2003; Simonson et al., 2022</u>). When performing a high MWL task, operators have to dedicate an important part of their cognitive resources to it, leaving few resources available to allocate to an incoming emergency alarm (<u>Wickens, 2008</u>; <u>2020</u>). In these conditions, operators may not be able to detect and properly process an incoming alarm (<u>Causse, Imbert et al., 2016; Causse et al., 2022</u>; <u>Giraudet et al., 2015</u>), a phenomenon referred to as *inattentional deafness* for auditory stimuli (e.g., <u>Dehais et al., 2019</u>; <u>Zhu et al., 2022</u>) and *inattentional blindness* for visual stimuli (e.g., <u>Lavie et al., 2014</u>; <u>Simons, 2000</u>; <u>White & O’Hare, 2022</u>). Perceptual load was also found to affect the detection and processing of incoming alarms in a similar way as MWL (<u>Jaquess et al., 2017</u>; <u>Lavie et al., 2014</u>; <u>MacDonald & Lavie, 2011</u>; <u>Molloy et al., 2015</u>; <u>Raveh & Lavie, 2015</u>). The deleterious effects of perceptual load and MWL are likely to combine in high MWL situations as the latter are usually characterized by elevated levels of visual (e.g., numerous displays to monitor) and auditory (e.g., auditory alerts, communications) perceptual load (<u>Causse et al., 2022</u>) with potential catastrophic consequences (see for instance Eastern Air Lines Flight 401: <u>NTSB, 1973</u>)

Even though an alarm has been successfully detected and processed by an operator, different factors, such as stress or MWL, can still impair the cognitive ability and psychomotor skills of operators and disrupt the response to the alarm (<u>Martin et al., 2015</u>, <u>2016</u>; <u>Schmidt et al., 2008</u>; <u>Whittemore & Woods, 2021</u>). The sudden and intense stress generated by emergency alarms can trigger inappropriate reactions in operators, such as freezing reactions (i.e., a defensive reaction consisting in a state of attentive immobility and body tension; <u>Roelofs, 2017</u>) that can slow down or even inhibit operators’ response to the alarm (<u>DeCelles, 1991</u>; <u>Heaslip et al., 1991</u>; <u>Martin et al., 2012</u>; Air Canada Flight 189: <u>TSB, 1998</u>), or startle reactions (i.e., “a simple reflex action that generally commences with an eye blink and develops into an aversive movement away from the stimulus”; <u>Martin et al., 2015</u>, <u>2016</u>; Air France Flight 447: <u>BEA, 2012;</u> Colgan Air Flight 3407: <u>NTSB, 2010</u>). High levels of MWL (e.g., competing task) can also affect the execution of the response to emergency alarms (<u>Bustamente et al., 2007</u>; <u>Shaw et al., 2018</u>), with potential catastrophic consequences (<u>Kuchar, 2001</u>; <u>NTSB, 2000</u>).

### 1.3 Improving operators’ reactions with gesture alarms?

Because operators have limited attention and working memory resources (<u>Jaquess et al., 2017</u>; <u>Wickens, 2020</u>), they should be presented with concise and easy-to-understand information (<u>Endsley & Jones, 2004</u>) that can spread out the workload and allow optimal responses (<u>Bliss & Gilson, 1998</u>). More specifically, <u>Whittemore and Woods (2021)</u> recommend using proactive instruction alarms (e.g., the “Pull Up” of the GPSW alarm) in emergency situations, instead of simple awareness warnings (e.g., Stall alarm).

Speech alarms are not the only way to communicate proactive instructions to the operators. Videos of the action to be performed in response to a specific emergency could also be used. <u>Fabre and colleagues (2021)</u> have recently proposed that, because action videos both strongly attract attention (<u>Abrams & Christ, 2003</u>) and preactivate the motor cortex (<u>Rizzolatti et al., 1996</u>), they might (compared to simple written speech alarms) facilitate both the detection of emergencies and the initiation of the expected motor responses in operators (<u>Brucker et al., 2015</u>; <u>Iacoboni et al., 1999</u>; <u>Jeannerod, 1994</u>; <u>Meltzoff & Prinz, 2002</u>).

To test this assumption, <u>Fabre and colleagues (2021)</u> conducted a two-experiment study investigating both 1) the response performances to a “Pull Up" hand action video (hereinafter referred to as gesture alarm) and the current GPWS written inscription and 2) the associated brain electrophysiological responses to these two types of stimuli. Faster reaction times and greater mu and beta desynchronizations were found in response to gesture alarms than to the current GPWS written inscription, suggesting that the faster reaction to gesture alarms might result from a greater motor preparation and pre-activation of “Pull Up” recovery maneuver (<u>Fadiga et al., 1995</u>). No difference in response accuracy was found between the two types of alarms. The authors attributed this result to the fact that participants always had to execute the same action (i.e., Pull Up) in response to the two types of alarms, which might have triggered automatic responses and erased a potential difference in accuracy.

Emergency situations are usually associated with a high MWL, which is known to affect both the detection and the response to emergency alarms (<u>Simonson et al., 2022</u>). It is therefore essential to evaluate whether and to what extent high levels of MWL might affect the response to new emergency systems before they are implemented in operational environments. Even though the preliminary results of <u>Fabre and colleagues</u> are encouraging, the impact of MWL on operators’ response performance (i.e., accuracy and reaction times) to gesture alarms compared to written inscription alarms remains unknown.

### 1.4 The present study

The present study aimed at investigating the efficiency of gesture alarms (compared to written alarms) under variable levels of MWL. Participants had to perform an *alarm-response task* drawn from the alarms (i.e., Stall and GPWS alarms) and the expected responses of two aviation emergency situations (i.e., incoming CFIS or CFIT). Participants had to respond to two opposite *gesture alarms* – i.e., videos of a hand either pushing or pulling a joystick – and the corresponding *written alarms* – i.e., written inscriptions of either the verb *“push”* or the verb *“pull”*. Participants had to respond to the alarms while performing: 1) no other competing task (no MWL), 2) a competing mental calculation task generating a low level of MWL and 3) a high level of MWL. Based on the results of <u>Fabre et al. (2021)</u>, we predicted to observe faster reaction times to gesture alarms than to written alarms and no difference in response accuracy between the two types of alarms in the no MWL run. We also predicted that 1) increasing participants’ MWL would overall slowdown and negatively affect the accuracy of their responses to both types of alarms and that 2) participants would be faster and more accurate in responding to gesture alarms than to the written alarms as the level of MWL increased.

The present study also aimed at investigating the event-related spectral perturbations (ERSPs) observed in response to the gesture alarms and the written alarms as a function of the level of MWL (e.g., <u>Behmer & Fournier, 2014</u>). The high temporal and frequential resolution of ERSPs allows to investigate the brain processes associated with the processing of motor stimuli (<u>Pfurtscheller & Da Silva,1999</u>). Mu and beta desynchronization were repeatedly found at parietal electrodes (i.e., C3 and C4) when participants had to observe and/or imitate an action (e.g., <u>Fox et al., 2016</u>; <u>Meyer et al., 2020</u>; <u>Pfurtscheller et al., 1997</u>; <u>Pineda, 2005</u>; <u>Quandt et al., 2012</u>), or read action language (e.g. <u>Moreno et al., 2013</u>; <u>Niccolai, et al., 2014</u>), reflecting the preparation and/or the execution of actions and sometimes interpreted as reflecting the activity of mirror neuron system. We predicted that, similarly to when participants only had to observe the alarms (<u>Fabre et al., 2021</u>), greater mu and beta power decrease would arise in response to gesture alarms than to written alarms, reflecting the greater facilitation of the response preparation and/or execution triggered by gesture alarms (compared to written alarms). We also made the assumption that these differences in mu and beta power between the two types of alarms would vary as a function of the level of MWL (e.g., <u>Behmer & Fournier, 2014</u>; <u>Jenson et al., 2019</u>; <u>Muthukumaraswamy & Singh, 2008</u>).

## 2 MATERIAL AND METHODS

### 2.1 Participants

Based on the literature in the field, reporting in general between 17 and 37 participants for this type of experimental design with EEG measures (<u>Mollo et al., 2016</u>; <u>Quandt et al., 2012</u>), 28 participants (*M*age = 26, *SD =* 8, age range: 20 48 years old; 12 females) took part in the present study. They were recruited thanks to paper ads that were disposed in various locations in the University campus. They were engineering students, PhD students or postdoctoral fellows. All had normal or corrected-to-normal vision and none of them reported a history of prior neurological disorder. All participants were informed of their rights and gave written informed consent for participation in the study.

### 2.2 Ethics Statement

This study was carried out in accordance with both the Declaration of Helsinki and the recommendations of the Research Ethics Committee of the University of Toulouse in France (CERNI no. 2018-107).

### 2.3 Material

#### 2.3.1 Alarms

Six different stimuli were presented to the participants. The stimuli of interest were two *written alarms* consisting in a red written inscription of the verb “PULL” or “PUSH” (respectively Figures 1.A and 1.C.) and two *gesture alarms* consisting in videos of a hand pulling or pushing a joystick (respectively Figures 1.D & 1.F).

**Figure 1.**
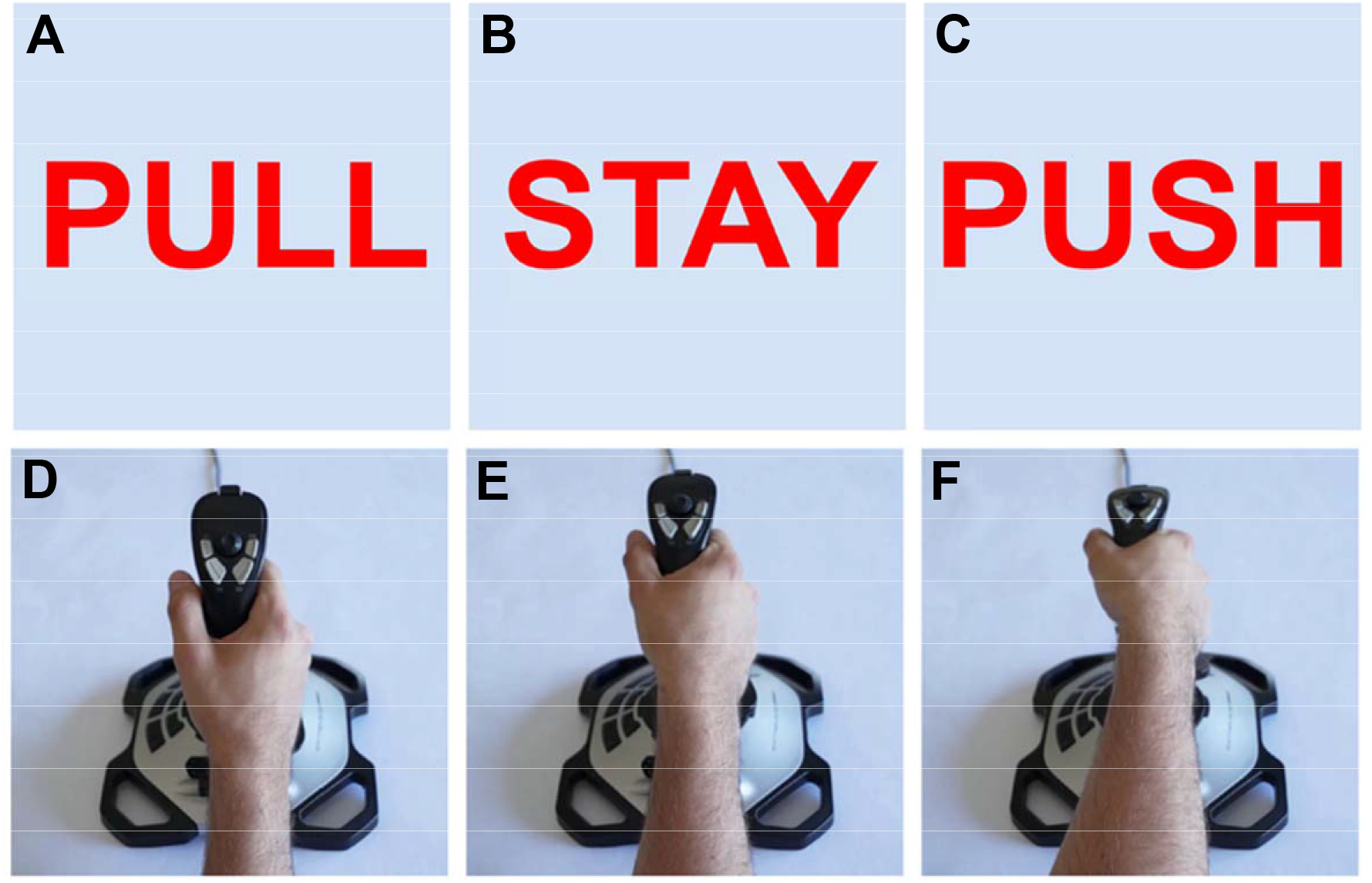
Illustration of the two (A) PULL and (C) PUSH written alarms and the (B) STAY written filler; and the corresponding (D) PULL and (F) PUSH gesture alarms and (E) gesture filler. For the written alarms and written filler, the verb was displayed twice for 1800ms followed by a grey background for 200ms each time. For the gesture alarms and the gesture filler, the video lasted 2000ms (the hand gesture lasted 1800ms and the hand remained still for 200ms at the end of the gesture) and was played twice.

To prevent automatic responses to the four alarms of interest, two filler alarms to which no response was expected were created. They consisted in a red written inscription of the verb “STAY” (i.e., written filler) and a video of a motionless hand on a joystick (i.e., gesture filler; respectively Figures 1.B. & 1.E.).

#### 2.3.2 Experimental apparatus

The experimental paradigm was presented using E-Prime 3 (Psychology Software Tools, Inc., Pittsburgh, PA) on a computer. Participants were seated in a chair and placed in front of a 19” monitor and had to respond to the alarms using a Logitech © Extreme 3D Pro joystick.

#### 2.3.3 Electroencephalogram (EEG)

The EEG signals were recorded and amplified with the Biosemi© system (Amsterdam, The Netherlands; https://www.biosemi.com/products.htm) using 64 Ag-AgCl active scalp electrodes (Fp1, AF7, AF3, F1, F3, F5, F7, FT7, FC5, FC3, FC1, C2, C3, C5, T7, TP7, CP5, CP3, CP1, P1, P3, P5, P7, P9, PO7, PO3, O1, Iz, Oz, POz, Pz, CPz, Fpz, Fp2, AF8, AF4, AFz, Fz, F2, F4, F6, F8, FT8, FC6, FC4, FC2, FCz, Cz, C2, C4, C6, T8, TP8, CP6, CP4, CP2, P2, P4, P6, P8, P10, PO8, PO4, O2) arranged according to the international 10-20 system. Two electrodes were placed on the mastoids. Four external electrodes were placed around the eyes to record the vertical (two electrodes) and horizontal (two electrodes) electro-oculogram (EOG). Two additional electrodes placed close to Cz, the Common Mode Sense [CMS] active electrode and the Driven Right Leg [DRL] passive electrode, were used to form the feedback loop that drives the average potential of the participant as close as possible to the AD-box reference potential (<u>Metting van Rijn, et al., 1990</u>). Skin-electrode contact, obtained using electro-conductive gel, was monitored, keeping voltage offset from the CMS below 20 mV for each measurement site as recommended by the manufacturer. All the signals were (DC) amplified and digitized continuously with a sampling rate of 512 Hz with an anti-aliasing filter with 3 dB point at 104 Hz (fifth order sinc filter), no high pass filtering was applied online. The triggers indicating the onsets of the alarms were recorded on additional digital channels.

### 2.4 Tasks

#### 2.4.1 Alarm response task

In the *alarm response task*, participants were presented with the four alarms of interest (i.e., push gesture, push written, pull gesture, pull written alarms) that were played 15 times each, and the two filler alarms (i.e., stay gesture, stay written alarms) that were played 30 times each. The alarms and the fillers were randomly presented to the participants for 4 seconds, with a variable inter-trial interval lasting between 4 and 6 seconds. Participants’ task consisted in 1) responding to the alarms of interest according to the action they described (i.e., pushing the joystick in response the push gesture and push written alarms and pulling the joystick in response the pull gesture and pull written alarms), and 2) remaining still when filler alarms were displayed (i.e., stay gesture and stay written alarm).

#### 2.4.2 Mental arithmetic task

The *mental arithmetic task* used in the present study was a subtraction task drawn from the Trier Social Stress Test (TSST; <u>Birkett, 2011</u>). Participants’ task consisted in repeatedly subtracting either 5 in the low MWL run or 13 in the high MWL run from a fixed starting number (1022). Participants had to orally state their response and continue their subtraction loop until 1) they reached the closest number to 0; or 2) they made a mistake. In the former case (i.e., participants made no mistake), they were asked to perform the same task a second time (and sometimes even a third time) from a different starting number (i.e., 1023 the second time and 1024 the third time) until the end of the run (see section 2.6. for further detail). In the latter case (i.e., participants made a mistake), the experimenter notified the participants of their mistake, and the participants were required to start all over again from the initial number (e.g., 1022).

### 2.5 Data measures

#### 2.5.1 Behavioral measurements

##### Alarm response task

Performances to the *alarm response task* were assessed in terms of accuracy (i.e., erroneous responses) and reaction times to correct responses. During the pre-tests, we found that the E-Prime default detection threshold of the responses was too sensitive for our experimental paradigm (i.e., micro-movements of the hand on the joystick were counted as responses). We determined that only considering the joystick inputs characterized by a vertical amplitude superior to 100 pixels above (i.e., pull) and below (i.e., push) the center of the screen (i.e., the default position of the joystick) allowed an optimal detection level of the responses on our 1920 x 1080 screen. As a result, only the joystick inputs superior to +/- 100 pixels on the vertical axis were considered to be responses to the alarms. Erroneous responses were computed as the sum of non-responses (i.e. when the participant did not respond to an alarm of interest) and inadequate responses (i.e., pulling in response to a push alarm and pushing in response to a pull alarm). Reaction times were defined as the time period between the onset of the alarm and the beginning of the participants’ response. The reaction times to erroneous responses and those above or below two standard deviations around the mean were eliminated (<u>Ratcliff, 1993</u>).

##### Mental arithmetic task

Performances to the modified TSST in the low MWL run and the high MWL run were assessed in terms of 1) number of mathematical operations, 2) number of mathematical errors and 3) error ratio, which was computed as the number of mathematical errors divided by the total number of operations.

#### 2.5.2 Subjective measures

At the end of each run, participants were asked to subjectively evaluate the difficulty of the run (i.e., performing both tasks irrespective of the type of alarm) they just completed by filling the six sub-scales of the NASA-TLX questionnaire which assess the perceived mental demand, the physical demand, temporal demand, performance, effort and frustration of participants (<u>Hart & Staveland, 1988</u>). Afterwards, they were asked to evaluate the extent to which they found it difficult to respond to 1) the written alarms and 2) the gesture alarms on two different 7-point Likert scales (1 -> *not difficult at all* to 7 -> *extremely difficult*).

#### 2.5.3 Brain electrophysiological measures

The Event-Related Spectral Perturbations (ERSPs) time-locked to the onset of written alarms and gesture alarms were investigated in all three levels of MWL (i.e., runs). To compute the latter, EEG data were analyzed using EEGLAB v.13.6.5b open-source software (<u>Delorme & Makeig, 2004</u>) on Matlab 2017a. First, EEG signals were band-pass filtered ([0.1-30] Hz) using a basic FIR filter. An independent component analysis was performed to isolate eye-blinks and movement-related artifacts that were removed from the signal (<u>Jung et al., 1998</u>). A visual inspection of the data was performed to reject residual artifacts intervals. Data were segmented into epochs from - 1000 to 2000 ms around the stimuli onset. Four participants were removed from the analysis due to the low quality of their EEG signal (less than 10 trials per condition). For the remaining 24 participants, epoched data were separated for each experimental condition according to the run (no MWL, low MWL and high MWL) and to the event type (gesture or written alarm). ERSPs were computed with Morlet wavelets as performed by EEGLAB, time-locked to the event display. They were computed with 30 frequencies in the 3-30 Hz window, three wavelet cycles at the lowest frequency, an increasing factor of 0.8 and 350 time-points in the - 441 to 1441 ms time-window (with 0 corresponding to the onset of the stimuli). Finally, as recommended ERSPs were baseline-corrected with a common ERSP baseline across factors and averaged separately for each condition across participants.

### 2.6 Procedure

After signing the inform consent form, participants were asked to take place in the experimental room. They sat in a comfortable seat in front of the 1920 x 1080 computer screen. Participants were asked to read the instructions of the experiment, while the experimenter placed a 64-electrode Biosemi EEG cap on their head.

Participants were told that they would have to perform the *alarm response task* under three different levels of MWL: once without any competing task (i.e., no MWL run), once simultaneously to a moderately demanding *mental arithmetic task* (i.e., low MWL run), and once simultaneously to a highly demanding *mental arithmetic task* (i.e., high MWL run). Participants were explained that they were expected 1) to respond as accurately and as fast as possible − when a response was expected − to different alarms in the *alarm response task* and 2) to make as many operations as possible without committing any calculation error in the *mental arithmetic task*. It was specified to the participants that the two tasks were of equal importance. In each run, participants were presented with 60 stimuli of interest and 60 fillers.

At the end of each run (lasting approximately 14 minutes), participants were asked to fill the NASA-TLX questionnaire (<u>Hart & Staveland, 1988</u>) and the two 7-point Likert scales evaluating the difficulty to react to respectively gesture and written alarms during the run. In order to prevent potential biases, the order of the runs was counterbalanced across participants and the instructions for the *mental arithmetic task* were provided to the participants just before the beginning of each run. The experimenter was sitting next to the participants during the three runs (but not when participants were filling the scales), evaluating the correctness of the calculations that were being made. The proximity of the experimenter aimed at both simulating the presence of a second operator and increasing participants’ MWL as a result of being watched (<u>Belletier et al., 2015</u>). At the end of the experiment, participants were debriefed by the experimenter and thanked for their participation in the study before they left the experimental room.

### 2.7 Data Analysis

#### 2.7.1 Behavioral data analysis

*Mental arithmetic task.* Three 2 [MWL (low MWL, high MWL)] paired t-tests were conducted on 1) the number of mathematical operations, 2) the number of mathematical errors and 3) the error ratio (*number of errors/number of operations*).

##### Alarm response task

As none of the variables was normally distributed (Shapiro-Wilk tests: *ps* < .05), a 3 x 2 [MWL (no MWL, low MWL, high MWL) x Alarm (written, gesture)] binary logistic regression was performed on the errors made in response to the alarms of interest (with 0 = correct response and 1 = erroneous response). A 3 x 2 [MWL (no MWL, low MWL, high MWL) x Alarm (written, gesture)] repeated-measures ANOVA analysis was conducted on the corrected log-transformed reaction times.

#### 2.7.2 Subjective data analysis

##### NASA-TLX

A 3 [MWL (no MWL, low MWL, high MWL)] one-way repeated measures ANOVA was conducted on the raw NASA-TLX data in order to assess the perceived task difficulty in the three experimental runs (<u>Hart, 2006</u>). The NASA-TLX sub-scales were also analyzed separately. As most variables were not normally distributed (Shapiro-Wilk tests: *ps* < .05), six 3 [MWL (no MWL, low MWL, high MWL)] one-factor Friedman’s ANOVAs were conducted on the NASA-TLX sub-scales. Wilcoxon signed-rank tests with a Bonferroni-adjusted alpha level of .017 (.05 / 3 comparisons) were conducted as post-hoc tests.

##### Difficulty to respond to the alarms

A 3 x 2 [MWL (no MWL, low MWL, high MWL) x Alarm (written, gesture)] ordinal logistic regression was performed on the ratings of difficulty to respond to the alarm from 1 (easy) to 7 (difficult), as none of the variables (with the exception of the difficulty to respond to the written alarms in the high MWL, *p* = .11) was normally distributed (Shapiro-Wilk tests: *ps* < .05). A manual stepwise analysis was performed to remove non-significant interactions from the model.

#### 2.7.3 EEG data analysis

Mean ERSPs were analyzed with permutation statistics (N = 2000 permutations) and a False Discovery Rate (FDR) correction for multiple comparisons with the run (i.e., no MWL, low MWL and high MWL) and event type (i.e., gesture alarm, written alarm) as within-subject factors. Decibel-corrected averages associated with power increase (positive values) or decrease (negative values) were then extracted for each frequency band i.e., delta [3 – 4[Hz, theta [4 – 8[Hz, mu [8 – 12[Hz, low beta [12 – 15[Hz and medium/high beta [15 – 30[Hz according to the time-window of significance identified by the permutation test at the C3 and C4 electrodes. ANOVA and Tukey post-hoc analyses were performed on the average power across participants in the frequency bands and time-windows in which the EEGLAB permutation test graphs revealed a significant main effect of MWL (see Figure 7.C & D). Paired t-test analyses were performed on the average power across participants in the frequency bands and time-windows in which the EEGLAB permutation test graphs revealed a significant main effect of the type of alarm (see Figure 7.C & D).

## 3 RESULTS

### 3.1 Behavioral results

#### 3.1.1 Mental arithmetic task

Participants made a significantly greater number of operations in the low MWL run [*t* (23) = 13.94, *p* < .001, CI95% (225.15, 303.60; *M* = 406.12, *SD* = 112.23] than in the high MWL run (*M* = 141.75, *SD* = 66.44; see Figure 2.A.). They also made a significantly greater number of mathematical errors in the high MWL run made [*t* (23) = - 3.31, *p* = .003, CI95% (- 6.10, - 1.41); *M* = 7.79, *SD* = 4.70] than in the low MWL run (*M* = 4.04, *SD* = 3.02; see Figure 2.B.). Finally, their error ratio was significantly greater in the high MWL run [*t* (23) = - 6.22, *p* < .001, CI95% (- .07, - .04); *M* = .06, *SD* = .04] than in the low MWL run (*M* = .01, *SD* =.01; see Figure 2.C.).

**Figure 2.**
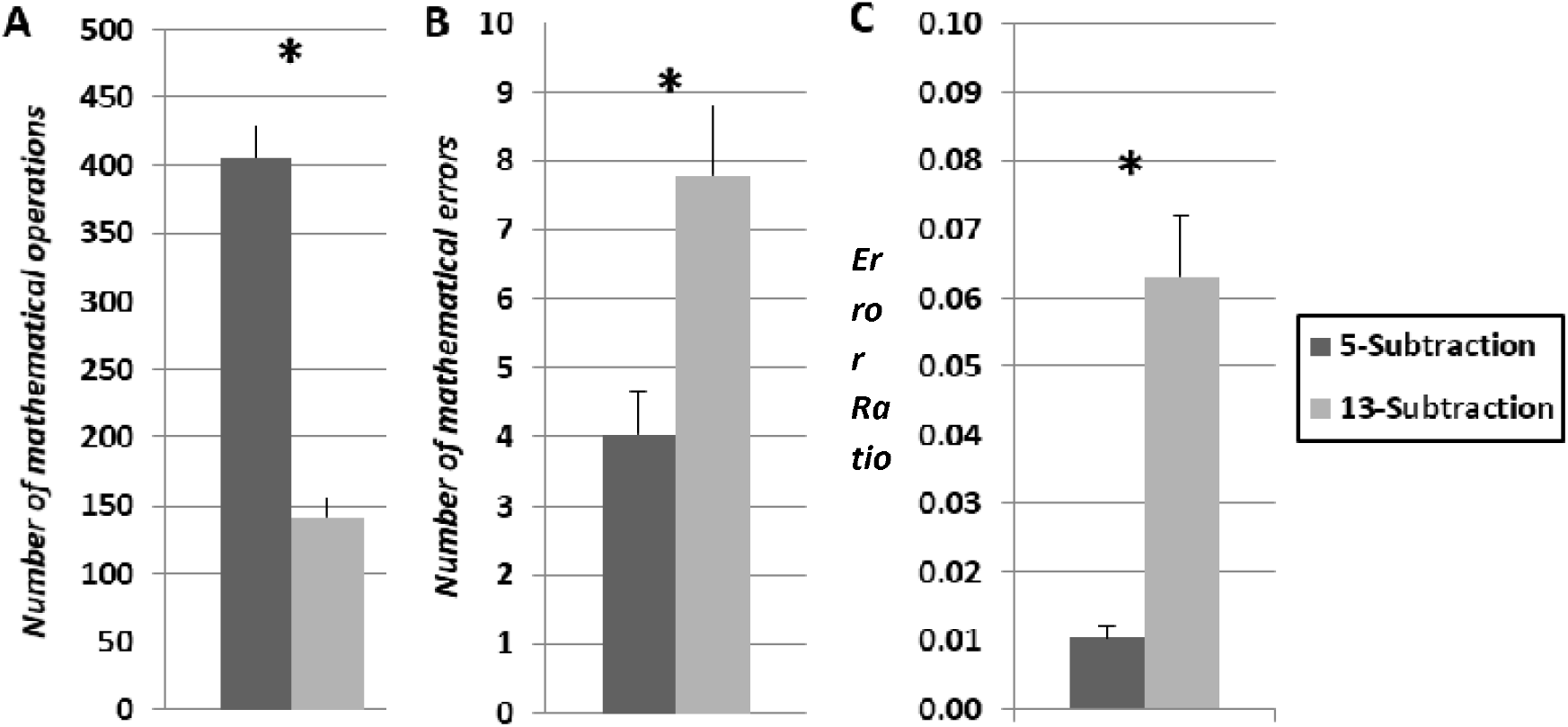
Illustration of (A) the number of mathematical operations, (B) the number of mathematical errors and (C) the error ratio in the low MWL run (dark grey) and in the high MWL run (light grey). * *p* < .01

#### 3.1.2 Alarm response task

##### Erroneous responses to the alarms

The analysis revealed a significant main effect of alarm [*B* (SE) = 2.833 (1.106), CI (95%) = (.666, 5.001), Wald χ*²* (1) = 6.565, *p* = .010; see Figure 3.A.], with gesture alarms (*M* = 3.50 %, *SD* = 6.08) predicting for a lower error rate than written alarms (*M* = 6.40 %, *SD* = 7.78). The analysis also revealed a significant main effect of MWL [no MWL: *B* (SE) = - 4.679 (1.270), CI (95%) = (- 7.169, - 2.189), Wald χ*²* (1) = 13.563, *p* < .001; low MWL*: B* (SE) = - 3.377 (1.362), CI (95%) = (- 6.048, - .707), Wald χ*²* (1) = 6.143, *p* = .013; with high MWL as dummy; see Figure 3.B.]. The high MWL run (*M* = 7.60 %, *SD* = 8.94) predicted for a lower accuracy than both the low MWL run (*M* = 4.34 %, *SD* = 5.50, *p* = .013) and the no MWL run (*M* = 2.91 %, *SD* = 5.64, *p* < .001), but no significant difference was found between the no MWL run and the low MWL run (*p* = .056).

**Figure 3.**
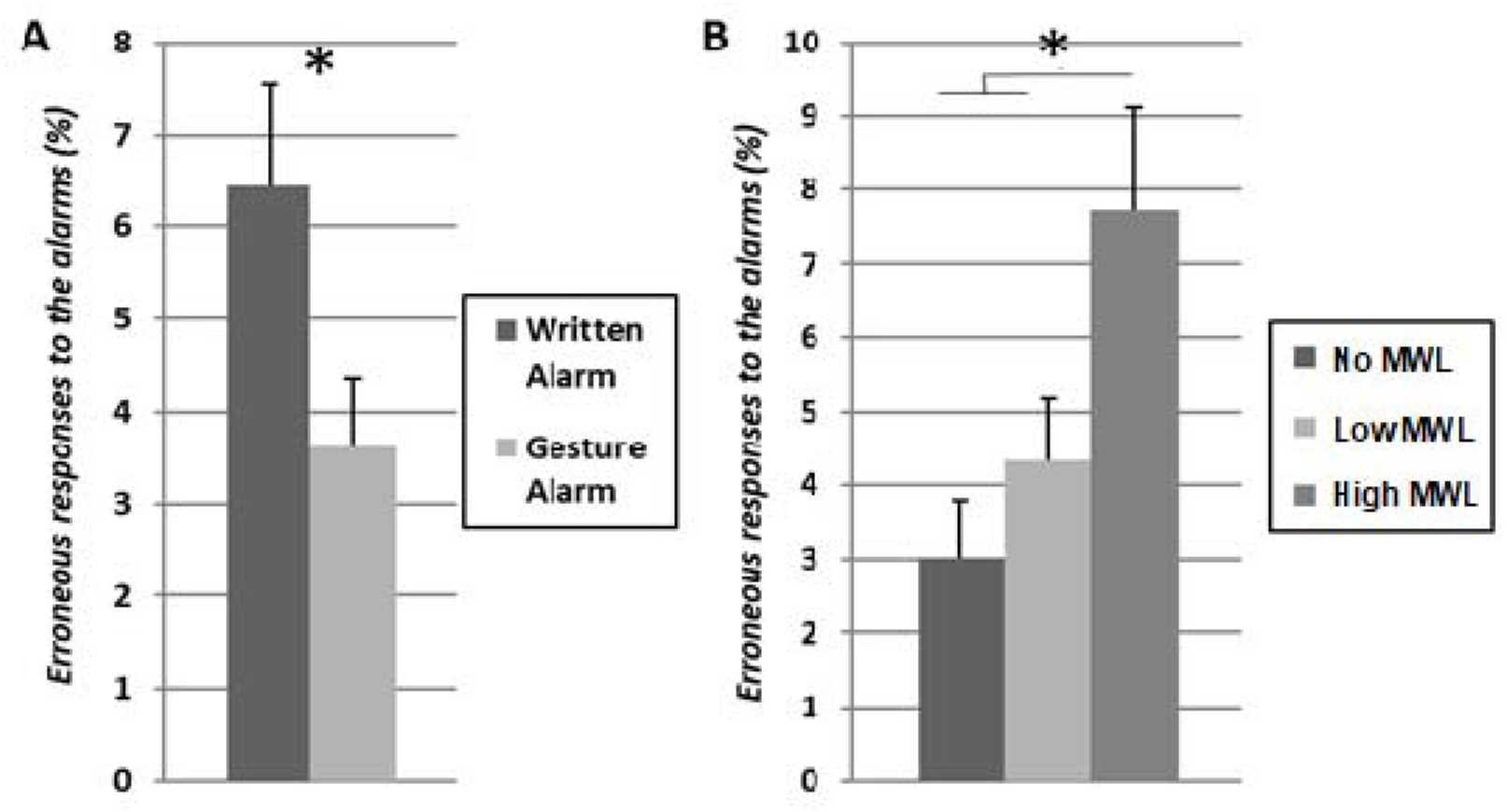
Illustration of the erroneous responses to the alarm (%) as a function of (A) the type of alarms with written alarms (dark grey) and gesture alarms (light grey); and (B) the level of MWL with no-MWL (dark grey), low MWL (light grey) and high MWL (mid grey). *: *p* < .05

##### Reaction times to the alarms

The analysis revealed a significant main effect of alarm [*F* (1, 23) = 34.56, *p* < .001, η*p²* = .60; see Figure 4.A.] with faster reaction times to gesture alarms (*M* = 930 ms, *SD* = 259) than to written alarms (*M* = 1140 ms, *SD* = 302). A significant main effect of MWL was also found [*F* (2, 46) = 46.75, *p* < .001, η*p²* = .67; Figure 4.B.], with longer reaction times in the high MWL run (*M* = 1228 ms, *SD* = 330, *p* < .001) and in the low MWL run (*M* = 1013 ms, *SD* = 236, *p* < .001) than in the no MWL run (*M* = 864 ms, *SD* = 200); and in the high MWL run than in the low MWL run (*p* < .001). The MWL x Alarm interaction did not reach significance [*F* (2, 46) = 1.06, *p* = .35, η*p²* = .04].

**Figure 4.**
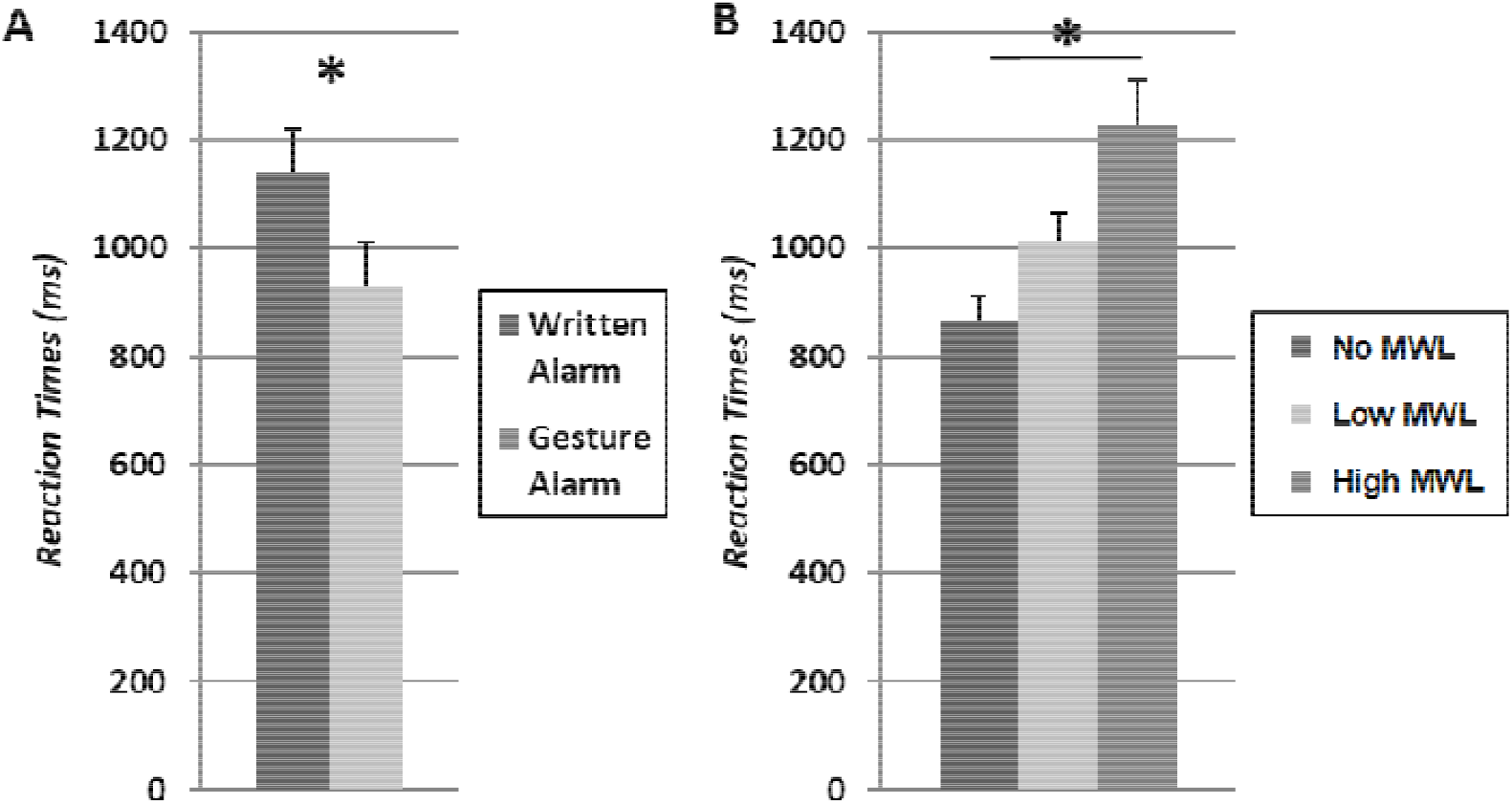
Illustration of the reaction times to the alarm (ms) as a function of (A) the type of alarms with writte n alarms (dark grey) and gesture alarms (light grey); and (B) the level of MWL with no MWL (dark grey), low MWL (light grey) and high MWL (mid grey). *: *p < .05*

### 3.2 Subjective results

#### 3.2.1 NASA-TLX questionnaire

The analysis revealed a significant main effect of MWL [*F* (2, 46) = 109.89, *p* < .001, η*p²* = 1.00; see Figure 5], with greater task difficulty perceived in high MWL run (*M* = 11.41, *SD* = 2.17, *ps* < .001) than in both the low MWL run (*M* = 9.89, *SD* = 2.28) and the no MWL run (*M* = 6.01, *SD* = 1.43); and in the low MWL run than in the no MWL run (*p* < .001).

**Figure 5.**
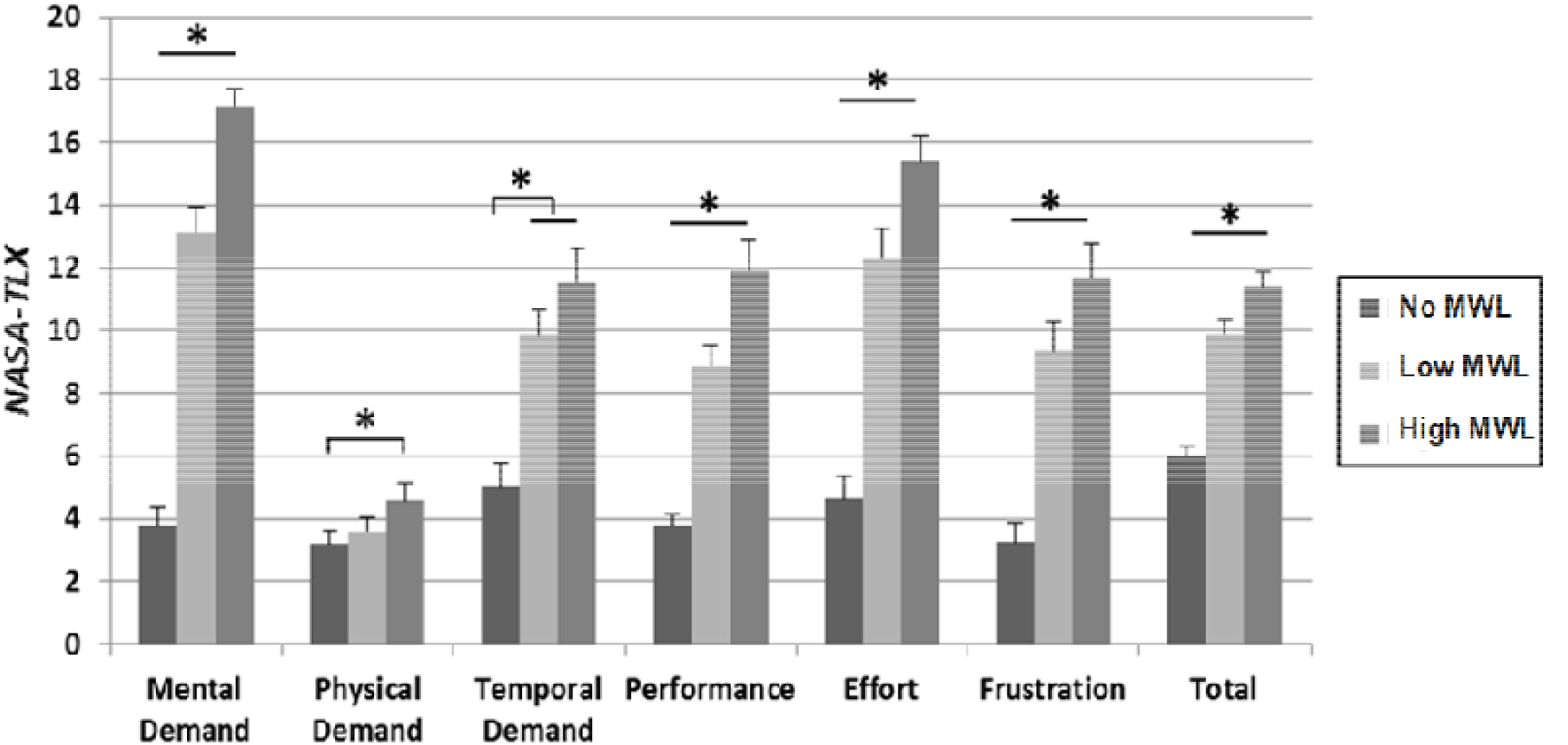
Illustration of the NASA-TLX ratings observed in the no MWL run (dark grey), the low MWL run (light grey) and the high MWL run (middle grey). *: *p* < .05

For the sake of clarity, the results of the statistical analysis conducted on the NASA-TLX subscales, the means and standard deviations are reported in Table 1.

**Table 1.** Summary of the means, standard deviations and statistical analysis performed on the ratings of NASA-TLX subscales.

| NASA-TLX | No MWL |  | Low MWL |  | High MWL |  | Friedman ANOVA |  | No vs. Low MWL |  |  | No vs. High MWL |  |  | Low vs. High MWL |  |  |
| --- | --- | --- | --- | --- | --- | --- | --- | --- | --- | --- | --- | --- | --- | --- | --- | --- | --- |
|  | *M* | *SD* | *M* | *SD* | *M* | *SD* | *X<sup>2</sup>F* (2) | *p* | *T* | *z* | *P* | *T* | *z* | *p* | *T* | *z* | *p* |
| **Mental Demand** | 3.79 | 2.81 | 13.13 | 4.09 | 17.17 | 2.53 | 45.66 | < **0.001\*** | 0.00 | 4.29 | < **0.001\*** | 0.00 | 4.29 | < **0.001\*** | 2.00 | 4.14 | < **0.001\*** |
| **Physical Demand** | 3.21 | 1.86 | 3.58 | 2.15 | 4.58 | 2.84 | 11.13 | **0.004\*** | 38.00 | 0.91 | 0.360 | 44.5 | 2.03 | **0.042\*** | 12.00 | 2.54 | **0.011\*** |
| **Temporal Demand** | 5.00 | 3.95 | 9.88 | 3.39 | 11.54 | 5.19 | 29.34 | < **0.001\*** | 17.50 | 3.79 | < **0.001\*** | 2.50 | 4.12 | < **0.001\*** | 52.50 | 2.18 | **0.029\*** |
| **Performance** | 3.79 | 1.74 | 8.87 | 3.33 | 11.92 | 4.65 | 40.06 | < **0.001\*** | 0.00 | 4.20 | < **0.001\*** | 0.00 | 4.20 | < **0.001\*** | 19.00 | 3.49 | < **0.001\*** |
| **Effort** | 4.62 | 3.46 | 12.29 | 4.76 | 15.37 | 4.13 | 41.93 | < **0.001\*** | 0.00 | 4.29 | < **0.001\*** | 0.00 | 4.20 | < **0.001\*** | 6.00 | 3.81 | < **0.001\*** |
| **Frustration** | 3.25 | 3.08 | 9.33 | 4.72 | 11.71 | 5.34 | 28.57 | < **0.001\*** | 8.00 | 3.95 | < **0.001\*** | 2.50 | 4.12 | < **0.001\*** | 49.00 | 2.31 | **0.021\*** |

#### 3.2.2 Difficulty to respond to the alarm

The analysis revealed a significant main effect of MWL [no MWL: *B* (SE) = - 1.625 (.244), CI (95%) = (- 2.103, - 1.147), Wald χ*²* (1) = 44.371, *p* < .001; low MWL*: B* (SE) = - .833 (.244), CI (95%) = (- 1.311, - .355), Wald χ*²* (1) = 11.67, *p* < .001; with high MWL as dummy; Figure 6A], with the no MWL run (*M* = 2.23, *SD* = 1.13) predicting for a lower difficulty to respond to the alarm than both the low MWL run (*M* = 3.02, *SD* = 1.49, *p* < .001) and the high MWL run (*M* = 3.85, *SD* = 1.48, *p* < .001); and the low MWL run than the high MWL run (*p* < .001). The analysis also revealed a significant main effect of type of alarm [*B* (SE) = 1.325 (.157), CI (95%) = (1.016, 1.634), Wald χ*²* (1) = 70.735, *p* < .001; Figure 6B] with gesture alarms (*M* = 2.37, *SD* = 1.41) predicting for a lower difficulty to respond than written alarms (*M* = 3.70, *SD* = 1.34).

**Figure 6.**
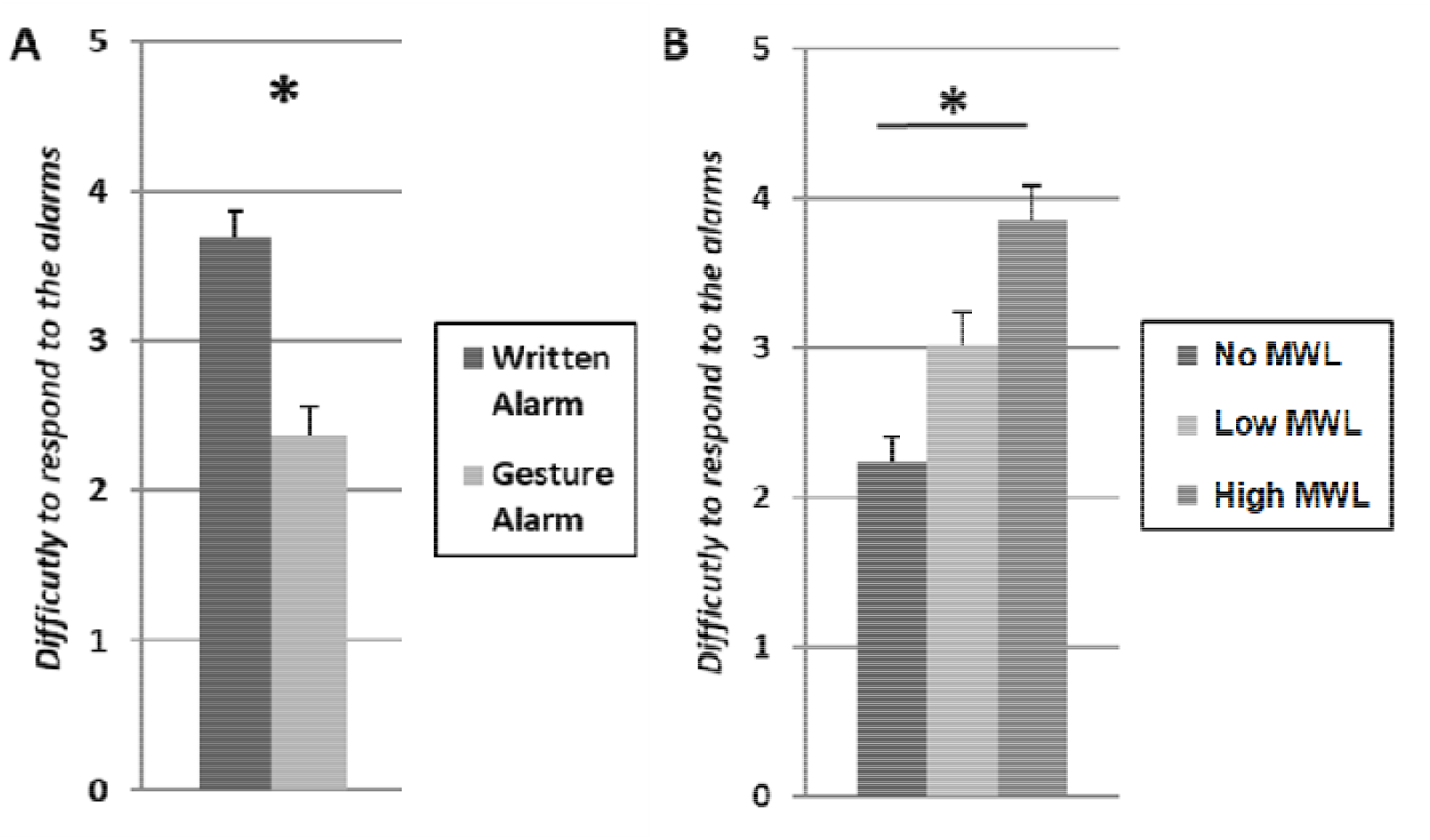
Illustration of the difficulty to respond to the alarms as a function of (A) the type of alarms with written alarms (dark grey) and gesture alarms (light grey); and (B) the level of MWL with no MWL (dark grey), low MWL (light grey) and high MWL (mid grey). *: *p* < .001.

### 3.3 Brain electrophysiological results

For the sake of clarity, the results of the effect of MWL and the type of alarm are reported respectively in Table 2 and Table 3, as well as displayed on Figure 7.

**Table 2.**
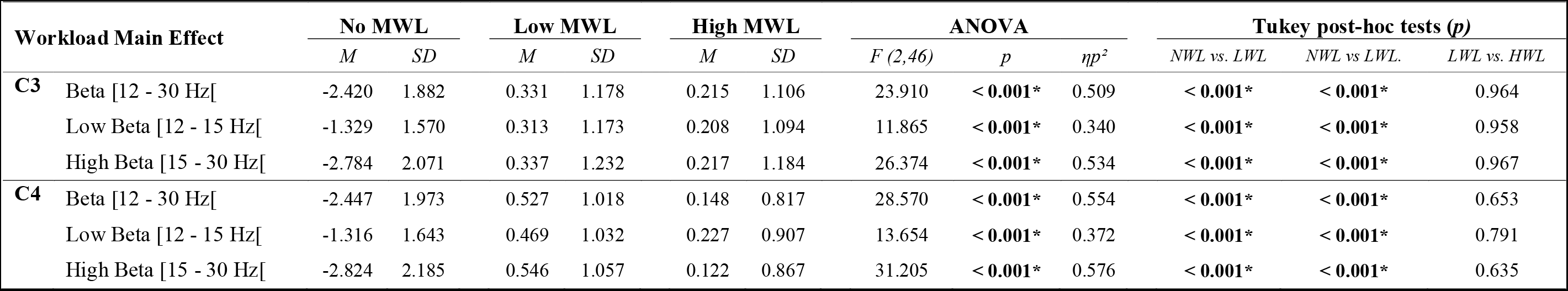
Summary of the means, standard deviations and statistical analysis performed on the total beta, low beta and high beta powers (dB) measured at C3 and C4 in response to the alarms in the no MWL (NWL), the low MWL (LWL) and the high MWL (HWL) runs in the 0 – 1441 ms time window after the onset of the alarms.

**Table 3.** Average power and standard deviations (dB) measured at C3 and C4 in response to both gesture and written alarms in the delta, theta [4 - 8 Hz[, mu [8 - 12 Hz[and low beta [12 - 15Hz[frequency bands in significant time windows. Time-windows were selected according to the results of the permutation tests with FDR correction for multiple comparison (see Methods section).

| Alarm Main Effect |  | Time window | Gesture Alarm |  | Written Alarm |  |
| --- | --- | --- | --- | --- | --- | --- |
|  |  |  | *M* | *SD* | *M* | *SD* |
| **C3** | Delta [3 - 4 Hz[ | 660 - 720 ms | 1.361 | 1.151 | 0.500 | 0.974 |
|  | Theta [4 - 8 Hz[ | 1000 -1440 ms | -0.893 | 1.151 | -0.159 | 1.026 |
|  | Mu [8 - 12 Hz[ | 1020 - 1350 ms | -1.523 | 1.463 | -0.867 | 1.568 |
|  | Low Beta [12 - 15Hz[ | 1170 - 1230 ms | -0.691 | 0.945 | 0.162 | 1.180 |
| **C4** | Delta [3 - 4 Hz[ | 590 - 820 ms | 1.511 | 1.156 | 0.430 | 0.940 |
|  | Theta [4 - 8 Hz[ | 970 - 1240 ms | -0.836 | 1.045 | -0.171 | 1.065 |
|  | Mu [8 - 12 Hz[ | 950 - 1100 ms | -1.850 | 1.391 | -1.040 | 1.640 |
|  | Low Beta [12 - 15Hz[ | / | / | / | / | / |

**Figure 7.**
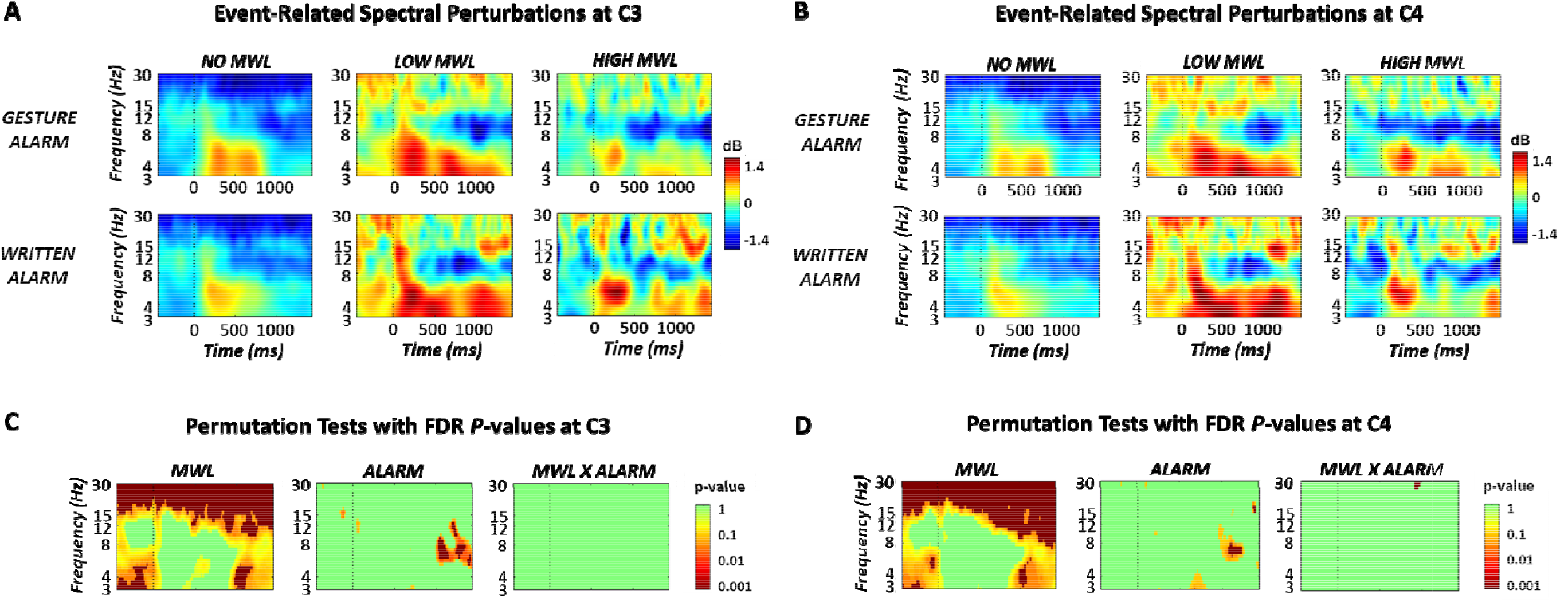
Event Related Spectral Perturbations (ERSPs) showing increases (positive values – red) and decreases (negative values – blue) in dB- corrected power in the three MWL levels (no MWL – left, low MWL – middle, and high MWL – right) for the two types of alarms (Gesture – top and written – bottom) at the C3 (A) and C4 (B) electrodes. Graphs C and D show results of the permutation tests for the MWL main effect (no MWL vs. low MWL vs. high MWL – right hand side top graph), the alarm main effect (gesture vs. written – bottom central graph) and the MWL x Alarm interaction (right hand side bottom graph). P-values are displayed with a log-scale from green (nonsignificant) to dark red (highly significant).

#### 3.3.1 Effect of the MWL

Permutation tests revealed a significant main effect of MWL at both the C3 and C4 electrodes in the low and high beta frequency bands (all *Fs* (2, 46) > 11; all *ps* <.001 and all η*p²* > .34; see Figure 7.C & D). Post-hoc analyses revealed that at both electrodes, the average power was significantly lower in the no MWL run than in both the low MWL and high MWL runs (all *ps* < .001; see Table 2). No difference was observed between the low MWL and high MWL conditions.

#### 3.3.2 Effect of the type of alarm

Permutation tests revealed a significant main effect of the type of alarm at both C3 and C4 in the delta, theta and mu frequency bands. Post-hoc analyses revealed an increase in the delta band, whereas we observed a decrease in both the theta and mu frequency bands for gesture compared to written alarms (all *ps* < .05; see Table 3 for the respective values and time windows). Finally, permutation tests also revealed a significant main effect of alarm in the low beta frequency band, but only at the C3 electrodes, with a lower activity for gesture compared to written alarms (*p* < .05; see Table 3 for the values and time window).

## 4 DISCUSSION

The aim of the present experiment was to investigate the efficiency of gesture alarms (compared to written inscriptions) under different levels of MWL at the subjective, behavioral and physiological levels. Participants performed an alarm response task concurrently to a mental subtraction task inducing several MWL levels.

### 4.1 Manipulation check of the mental arithmetic task

In line with literature, participants’ performance decreased (i.e., less mathematical operations, more calculation errors and higher error ratio) in the high MWL run compared to the low MWL run (<u>Birkett, 2011; Linares et al., 2020</u>). The analysis of the NASA-TLX ratings (<u>Hart, 2006</u>) also revealed that participants reported significantly higher levels of perceived task difficulty during the high MWL run than during both the low MWL and the no MWL runs; and during the low MWL run than the no MWL run. Taken together, these results serve as a manipulation check and confirm that the level of MWL significantly increased between the no, the low and the high MWL runs (<u>Feltman et al., 2022</u>; <u>Wickens, 2020</u>).

### 4.2 Impact of MWL on the response to the alarms

While participants took significantly longer and found it more difficult to respond to the alarms in the low MWL run than in the no MWL run, no difference in error rates to the alarm-response task was found between these two runs. These results suggest that even though the increase in MWL between the no MWL run and the low MWL run slowed participants’ reactions, enough attentional and cognitive resources were still available in the low MWL run to maintain a similar level of response accuracy as in the no MWL run (<u>De Sanctis et al., 2014</u>; <u>Wickens, 2008</u>). In the high MWL run, participants took significantly longer, made more erroneous responses, and found it harder to respond to both types of alarm than in both the no MWL run and the low MWL run. First, the general decrease in performance to both the *mental arithmetic task* (i.e., number of operations and number of errors) and the *alarm- response task* (i.e., both accuracy and speed; in line with literature; e.g., <u>Causse et al., 2022; Giraudet et al., 2015</u>) show that participants followed the instructions they were given, that is to consider the two tasks as of equal importance and not prioritize one over the other. Second, these results also suggest that the reserve of attentional and cognitive resources was not sufficient to enable the completion of the *alarm-response task* in the high MWL run with the same level of performance, as in the no MWL and low MWL runs (<u>Jaquess et al., 2017; Wickens, 2008</u>; <u>2020</u>), resulting in a significant drop in response accuracy and longer reactions times (in line with <u>De Sanctis et al., 2014</u>; <u>Diekfuss et al., 2017; Shaw et al., 2018</u>).

At the brain electrophysiological level, greater beta activity was found in response to the alarms in general in the low MWL and high MWL runs than in the no MWL run, along the whole epoch. Greater beta activity was found to reflect increased task demand/difficulty, (mental) workload and/or (mental) fatigue (e.g., <u>Kumar & Kumar, 2016</u>; <u>Michels et al., 2010</u>; <u>Morales et al., 2019</u>; <u>Okogbaa et al., 1994</u>; <u>Rietschel et al., 2012</u>). Therefore, the greater beta activity observed in response to the alarms in both the low and high MWL runs may reflect the increase in task demand triggered by the mental arithmetic task. The fact that the difference in beta activity was found on the whole epoch (even before the alarms were presented) suggests that this increase in task demand might have been sustained along the whole run, confirming once again the efficiency of this mental arithmetic task to trigger MWL in participants.

### 4.3 Gesture alarms versus written alarms

Participants were found to be both faster and more accurate when responding to the gesture alarms than to the written alarms (in line with <u>Fabre et al., 2021</u>), regardless of the level of MWL. They also reported that they found it easier to respond to the gesture alarms than to the written alarms, once again regardless of the level of MWL. Taken together, these results suggested that gesture alarms might be more efficient to trigger fast and accurate responses than simple written instructions, even in high MWL situations.

In line with our predictions and the previous results of <u>Fabre and colleagues (2021</u>), greater mu and low beta power decreases were found in response to gesture alarms than to written alarms. Mu and beta power desynchronizations are known to reflect the preparation and the execution of actions (e.g., <u>Brinkman et al., 2016</u>; <u>Fox et al., 2016</u>; <u>Meyer et al., 2020</u>; <u>Pfurtscheller et al., 1997</u>; <u>Pineda, 2005</u>; <u>Quandt et al., 2012</u>). The results of the present study indicate that, compared to written alarms, gesture alarms might facilitate the preparation and/or the execution of the response. A greater power decrease in the mu frequency band was found for gesture alarms than for written alarms in time-window of the response to the alarms (i.e., on average 930ms for the gesture alarms and 1140ms for the written alarms). These results suggest that the greater decrease in power observed in response to gesture alarms is more likely to reflect the facilitation of the action execution, rather than the facilitation of its preparation. However, since we were not able to investigate the signal associated with the onset of the response observed for both gesture and written alarms (i.e., no triggers were sent when participants performed the action), we cannot conclude with certainty on this point.

The results also suggest a greater high delta (3-4 Hz) synchronizations in response to the gesture alarms than to the written alarms. Increased delta activities were found to underpin action selection and/or execution in cognitively demanding situations (<u>Beste et al., 2012</u>) and to be correlated to performance at action selection (<u>Hamel-Thibault et al., 2018</u>). In the present study, the greater high delta activities observed in response to gesture alarms than to written alarms occurring before the execution of the action could reflect the facilitation of the action selection in response to gesture alarms. However, this result should be taken with caution because the present study was not designed to investigate the delta rhythm. First, the time-frequency tradeoff intrinsic to the definition of Morlet wavelets did not create a sufficient precision in the frequency domain to analyze very low frequency activities with three cycles. Second, this small number of cycles which is required to reach a sufficient time-domain accuracy at low frequencies induces a frequency leakage from close-by frequencies when transposed to the frequency domain (<u>Cohen, 2014</u>). Consequently, the delta power measured by the EEG could have been polluted by surrounding (e.g., theta) frequency activities. Further research is therefore necessary to confirm this potential difference in delta synchronization between gesture and written alarms.

### 4.4 Applications, limitations and future works

Compared to written instruction alarms, gesture alarms were found to facilitate the initiation of the expected response and to trigger faster and more accurate responses under all three MWL levels. The results of the present study therefore suggest that gesture alarms could be a suitable solution to improve both the visual modality of emergency alarms and the operators’ performances, even under high levels of MWL (<u>Bliss & Gilson, 1998; Endsley & Jones, 2004</u>).

In the context of aviation, gesture alarms could be used to improve the GPWS alarm, and more importantly the Stall alarm (<u>Whittemore & Woods, 2021</u>). While the maneuver that has to be performed in reaction to the Stall alarm is both stressful and counter-intuitive for pilots (<u>BEA, 2012</u>; <u>IATA, 2018a</u>; <u>NTSB, 2010</u>), the current Stall alarm consists in a simple awareness warning (i.e., only indicating the dangerous deviation of a parameter; <u>Ewbank, et al., 2016</u>). Replacing the visual modality of the Stall alarm by a gesture alarm (i.e., hand video pushing on the stick) might significantly enhance its design and improve pilots’ response performance to it.

The implementation of gesture alarm systems might also help improve the reactions of operators in many other fields where fast and accurate responses are needed. Gesture alarms may be particularly adapted to nuclear powerplants and transportation in general (i.e., air, rail, or road transport), to improve emergency shutdown or collision avoidance maneuvers for instance. They might also be used in the medical field (<u>Alizezaee et al., 2017</u>; <u>Edworthy, 2013</u>).

However, further research is needed before gesture alarms could be implemented in real operational environments. First, while most alarm systems are multimodal, we focused exclusively on the visual component of the alarms. While, it is very likely that adding an auditory component will improve even more the operators’ response performances to gesture alarms (<u>Alirezaee et al., 2017</u>; <u>Child & Wendt, 1938</u>; <u>Hughes et al., 1994</u>; <u>Liu, 2001</u>), it remains to be demonstrated. More specifically, new studies should be conducted to determine which auditory component among symbolic sounds, simple auditory awareness warnings (e.g., “stall”), auditory pro-active instructions (e.g., “push down”) or a combination of these auditory alarms (<u>Whittemore & Woods, 2021)</u> would maximize the efficiency of gesture alarms. Second, the fact that gesture alarms were not tested on real-life operators in their usual work environment is another important limitation of the present study. Before gesture alarms could be implemented in work environments, they should be thoroughly tested on trained operators in the different operational contexts in which they could be implemented.

## Conclusion

The behavioral results of the present study demonstrate that compared to written instructions gesture alarms trigger faster and more accurate responses, regardless of the operators’ level of MWL. The brain electrophysiological results suggest that this greater efficiency might be due to the fact that (compared to written alarms) gesture alarms facilitate the execution (and also possibly the preparation) of the associated action. While further research is necessary to confirm the greater efficiency of gesture alarms in real operational environments, our study provides a step forward towards the implementation of such gesture-based warnings to improve operators’ performances in emergency situations.

